# Optimization of TMS target engagement: novel evidence based on combined TMS–EEG and dMRI tractography of brain circuitry

**DOI:** 10.1101/2024.08.19.607213

**Authors:** Pantelis Lioumis, Timo Roine, Ida Granö, Dogu Baran Aydogan, Elena Ukharova, Victor H. Souza, Dubravko Kičić, Risto J. Ilmoniemi, Nikos Makris

## Abstract

Neuromodulation is based on the principle that brain stimulation produces plastic changes in cerebral circuitry. Given the intersubject structural and functional variability, neuromodulation has a personalized effect in the brain. Moreover, because of cerebral dominance and interhemispheric functional and structural differences in the same individual, the characterization of specific brain circuitries involved is currently not feasible. This notion is extremely important for neuromodulation treatments applied in neuropsychiatry. Specifically, the efficacy of the neuromodulation treatments is critically dependent on the anatomical precision of the brain target and the circuitry which has been affected. However, a complete understanding of how the brain behaves under stimulation needs a combined characterization of its neurophysiological response. This can be achieved by TMS–EEG guided by current multimodal neuroimaging techniques in real time. Herein, we present novel data based on dMRI tractography-guided TMS–EEG on one healthy young adult volunteer.

## Introduction

The localization of cerebral cortical cytoarchitectonics is a critical concept and has been emphasized since late 1880’s by prominent neuroanatomists such as Brodman and Vogt (Brodmann’s Localisation in the Cerebral Cortex, 2005; Vogt and Vogt, 1919), as well as later by Von Economo, Bailey and Von Bonin (Bailey and Von Bonin, 1957; Zellaufbau der Grosshirnrinde des Menschen, n.d.), and recently by the neuroimaging community (Human Connectome Project; Glasser et al., 2016; Rushmore et al., 2022; Van Essen et al., 2019, 2012; Van Essen and Glasser, 2018). Cortical localization is associated with the structural circuitry of a given area. Adjacent areas may have vastly different connectivity matrices. This is related to differences in their cytoarchitectural structure; functional and epigenetic factors may have contributed to the differences as well, making connectivity highly individualized. These considerations are key concepts for neuromodulation, and more specifically for target engagement during brain stimulation. Given that neuromodulation is mechanistically related to Hebbian learning enabled by neuroplasticity, target engagement needs to be defined not only in terms of stimulated cortical area but also, importantly, in terms of brain circuitry associated with that area. Equally important is the characterization of the neurophysiological signature of the stimulated area, which should be combined with the specific underlying structural neuroanatomical background (cortical area and circuitry). Such an approach has not been achieved to date; therefore, using existing advanced neuroimaging and neurostimulation technologies, we demonstrate the feasibility to overcome this challenge, and to achieve stimulation specificity in target engagement in one healthy subject. To our knowledge, this is the first demonstration, the proof-of-principle of this approach.

## Methods

One subject (female, 21 years old) participated in the experiment. We first performed the MRI and then, a week after, the TMS–EEG acquisition.

### MRI acquisition

MRI data were acquired with a Siemens 3T Skyra. Diffusion MRI data were acquired with a state-of-the-art multi-shell sequence with the following parameters: a resolution of 2.0 × 2.0 × 2.0 mm^3^, simultaneous multislice (SMS) factor of 3, repetition time (TR) of 5000 ms, echo time (TE) of 100 ms, and 101 gradient orientations distributed in multiple diffusion-weighting shells of 900, 1600, and 2500 s/mm^2^. We acquired ten images with b=0 s/mm^2^ and three images with reverse-phase encoding for correcting echo planar imaging (EPI) distortions (Andersson et al., 2003).

T1-(with and without fat suppression) and T2-weighted MRI data were acquired with a resolution of 1.0 × 1.0 × 1.0 mm^3^. In addition, three T1-weighted images were acquired with a resolution of 0.8 × 0.8 × 0.8 mm^3^. Fat-suppressed T1-images were used for the generation of high-quality headmodels for realistic E-field modelling performed in SimNIBS (Thielscher et al., 2015). The MRI acquisition took one hour.

### T1-weighted image analysis (incl. parcellation)

The three regions of interest (pre-SMA proper, pre-SMA posterior, and SMA) were segmented manually based on anatomical criteria and visually recognizable landmarks on T1-weighted images. FreeSurfer was used for preprocessing and cortical parcellation of the T1-weighted images (Destrieux et al., 2010; Fischl, 2012). In addition, a Bayesian model was used to segment subcortical gray matter structures (Patenaude et al., 2011).

### Diffusion MRI data analysis

Preprocessing of diffusion MRI data included denoising (Veraart et al., 2016) followed by a correction for subject motion (Leemans and Jones, 2009), distortions caused by eddy currents (Andersson and Sotiropoulos, 2016), magnetic susceptibility (Andersson et al., 2003), and Gibbs ringing (Kellner et al., 2016). B1 field inhomogeneity correction was performed in ANTs (Tustison et al., 2010). Statistical parametric mapping was used to rigidly coregister T1-weighted and distortion-corrected diffusion MRI data (Penny et al., 2011).

Structural brain connectivity was measured from diffusion MRI data with whole-brain tractography (Tuch et al., 2002) performed with constrained spherical deconvolution (CSD) streamlines tractography (Tournier et al., 2019, 2007). With CSD, complex fiber configurations can be reliably estimated and thus, tractography through regions with crossing fibers can be performed (Jeurissen et al., 2013; Tournier et al., 2007). Multi-shell multi-tissue CSD (Roine et al., 2015; Dhollander et al., 2019; Jeurissen et al., 2014) was used to reconstruct 100 million streamlines by seeding from the white matter–gray matter interface using the probabilistic 2^nd^ order integration (iFOD2) algorithm in MRtrix3 (Tournier et al., 2019, 2007). Anatomically constrained tractography was used to improve the anatomical feasibility of the streamlines (i.e., that they begin from gray matter and pass through white matter until gray matter or the spinal cord is reached) (Smith et al., 2012). Regions of interest defined based on the coregistered T1-weighted images were used to extract structural connections specific to them from the whole-brain tractogram.

### TMS–EEG acquisition

The subject was seated in a comfortable chair and was instructed to relax and fixate a black cross 3 meters away during recordings. The experiment was carried out with a Nexstim-neuronavigated TMS system (NBT 5.0, Nexstim Plc., Finland) with a 70-mm radius cooled figure-of-eight coil and individual T1-weighted magnetic resonance images. The EEG signals were recorded with a BrainAMP EEG-system (Brain Products, Gmbh, Germany) with a 62-channels Easy cap with passive Ag/AgCl-sintered electrodes. The EEG signals were low-pass filtered at 1,000 Hz and sampled at 5,000 Hz.

The EEG electrodes were prepared by scraping the skin under each electrode with an abrasive paste (OneStep AbrasivPlus, H + H Medical Devices, Germany), after which each electrode was filled with a conductive gel (Electro-Gel, ECI, Netherlands), to keep the impedances below 5 kΩ throughout the whole experiment. The ground and reference electrodes were placed on the right zygomatic bone and mastoid, respectively. Electromyography was recorded with Nexstim EMG; the electrodes were placed in a belly-tendon montage on the right abductor pollicis brevis (APB) muscle.

During the recordings, active noise masking consisting of white noise with mixed-in click sounds (Russo et al., 2022; https://github.com/iTCf/TAAC) was played to the subject through in-ear earphones (ER3C Insert Earphones, Etymotic Research Inc., United States). The noise level was adjusted before the experiment until the subject was unable to hear the click at 80% maximum stimulator output (MSO) with the coil held 10 cm above the vertex. The absence of auditory evoked potentials was confirmed by visual inspection of 30 trials.

The APB hotspot was identified as the coil location and orientation that produced maximal and consistent motor-evoked potentials (MEPs). The resting motor threshold (rMT) was estimated with Nexstim’s built-in algorithm (Awiszus, 2003). The rMT was 61% MSO.

A total of seven recordings along the midline were performed (pre-SMA to M1) according to the parcellations. First, the most anterior target was mapped for a coil location, orientation and intensity that produced artefact-free and TEPs with early peak-to-peak amplitudes (< 50 ms) of 6–10 µV, resulting in a stimulation intensity of 71% MSO. Next, for each target in the parcellation (see Fig. 1), a good coil location was determined based on averaging 20 TEPs, after which recordings consisting of 100 pulses were measured from each target. The same coil orientation (lateral-medial) and intensity were used for all targets.

**Figure 1.**
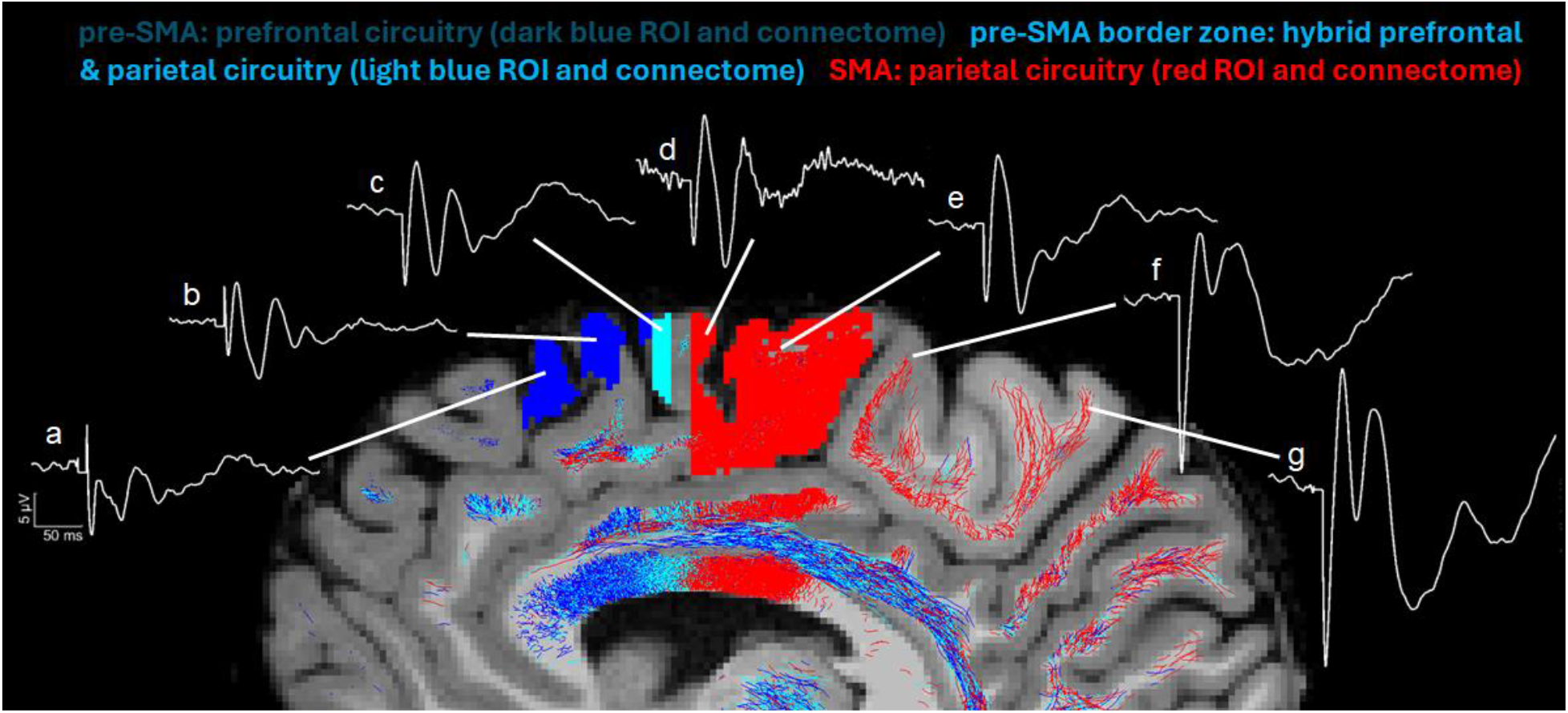
Neurophysiological signatures of distinct anatomical areas, in on healthy subject, based on TMS-EEG evoked responses (TEPs). Averaged TEPs (see supplementary material for details) have been recorded by a) pre-SMA superior, b) pre-SMA superior posterior, c) SMA anterior, d) SMA proper anterior, e) SMA proper posterior, f) leg premotor cortex, and g) leg primary motor cortex.

### TMS–EEG analysis

The TMS–EEG data was pre-processed in MATLAB R2024a with a script based on eeglab2021.0 (Delorme and Makeig, 2004) and the TESA package (Rogasch et al., 2017). Epochs were generated by selecting −1.5…1.5 s around each TMS pulse, after which baseline correction was applied (baseline time period: −100…−5 ms) by subtracting the baseline average from the signals. The TMS pulse artefact was removed by cutting and interpolating −2…6 ms around the TMS pulse with a cubic interpolation. Channels and trials were visually inspected, and those with high noise were removed (1 channel and 8 trials per recording were removed on average). Then, independent component analysis was applied to remove ocular artefacts from the data. Another baseline correction was applied, and the SOUND (lambda = 0.01) and SSP–SIR algorithms (0–10 ms) (Mutanen et al., 2018, 2016) were utilized to remove remaining noise and muscle artefacts. A bandpass filter from 0 to 200 Hz and a notch filter from 48 to 52 Hz were applied to remove high-frequency and line noise. Finally, the terminals were snipped to −1…1 s around the TMS pulse. For each recording, the channel with the largest range in the time window 15…50 ms after the TMS pulse was chosen for visualization (pre-SMA recordings: F1; SMA recordings: FC1; premotor and M1: CP1).

From the TEPs, we extracted their deflection peak amplitudes and latencies as shown in Table 1, but also the dominant frequencies. The dominant frequencies were calculated following the approach of (Rosanova et al., 2009a). For each channel, the event-related spectral perturbation (ERSP) was calculated with EEGLAB (Delorme and Makeig, 2004). In other words, for each channel, the power spectrum was calculated (window span: 3 to 0.8 wavelet cycles) over all trials, after which the trials were averaged and normalized by subtracting the mean baseline. A bootstrap approach was utilized to only consider significant activation with respect to the baseline (alpha = 0.01). The global ERSP was calculated by averaging the ERSPs over all channels. To estimate the dominant frequency, the ERSP was summed over the time interval 20…100 ms, after which the strongest frequency was selected within the 8…50 Hz interval.

**Table 1.** Peak amplitudes and latencies of all the averaged TEPs from each target, and also the dominant frequency in which each cortical site was perturbed (see below Fig. 1).

| Target/<br>peaks | I | II | III | IV | V | VI | VII | Dominant<br>frequency |
| --- | --- | --- | --- | --- | --- | --- | --- | --- |
| a | -6.5 μV<br>12 ms | -0.9 μV<br>19 ms | -5 μV<br>44 ms | 0.5 μV<br>58 ms | -3.2 μV<br>82 ms | -1.8 μV<br>93 ms | 1.7 μV<br>156 ms | 35 Hz |
| b | -3.1 μV<br>11 ms | 4.5 μV<br>20 ms | -5.4 μV<br>43 ms | 2.8 μV<br>60 ms | -2.1 μV<br>82 ms | -0.1 μV<br>93 ms | 0.8 μV<br>150 ms | 34 Hz |
| c | -7.4 μV<br>10 ms | 5.3 μV<br>21 ms | -6.3 μV<br>42 ms | 2.5 μV<br>58 ms | -3.9 μV<br>82 ms | -2.3 μV<br>94 ms | 3.0 μV<br>155 ms | 35 Hz |
| d | -6.7 μV<br>10 ms | 6.8 μV<br>21 ms | -9.0 μV<br>43 ms | 4.4 μV<br>62 ms | -2.8 μV<br>102 ms | | 3.0 μV<br>143 ms | 29 Hz |
| e | -7.9 μV<br>9 ms | 6.9 μV<br>23 ms | -9.2 μV<br>45 ms | -2.8 μV<br>68 ms | -3.7 μV<br>78 ms | -1.4 μV<br>94 ms | 2.9 μV<br>150 ms | 43 Hz |
| f | -18 μV<br>9 ms | 6.4 μV<br>28 ms | 0.1 μV<br>41 ms | 5.3 μV<br>68 ms | -7.1<br>120 ms | | | 39 Hz |
| g | -18 μV<br>10 ms | 12.4 μV<br>27 ms | -3.1 μV<br>43 ms | 8.3 μV<br>61 ms | -12.1<br>104 ms | | -1.2 μV<br>154 ms | 16 Hz |

## Results

Our results are shown in **Fig. 1**. We recorded TEPs that, although acquired with the same parameters (intensity and coil orientation), were different from each other in terms of peak amplitudes, but also in the peak timings, as can be observed below. It is also apparent that Peaks II and I (see Table 1), which represent reactivity, are considerably different between c and b, f and e, and g and f, as well as the dominant frequencies for e, f and g. Large amplitude differences can be seen between peaks VI and VII, whereas in some recordings, these deflections were not evoked.

## Discussion

Our results from the detailed stimulation of pre-SMA and SMA in one healthy subject demonstrate apparent and big differences in the features of the averaged TMS evoked signal. These features, early and late peak amplitudes, represent cortical reactivity and connectivity characteristics, respectively (Ilmoniemi et al., 1997; Massimini et al., 2005). In addition, the dominant frequency changes may imply that TMS–EEG can be a tool to identify adjacent changes in the cytoarchitectonics, as it has been earlier suggested for areas more distantly located from each other (Rosanova et al., 2009).

### From non-specific stimulation targeting to anatomically- and electrophysiologically-specified target engagement

A major question about TMS interventions is whether the stimulated structural circuitry is specific or not. Current neuroimaging technologies and analysis methods have not been applied efficiently enough to guarantee a precise characterization of cortical targets and their associated circuitry engagement specificity. Cortical anatomical connections are precise and architectonically specific. That is, each cortical field has a specific structural signature in terms of its laminar architecture and the fiber connections related to its different layers (Mesulam and Mesulam, 2000; Pandya et al., 2015; Pandya and Yeterian, 1985). This level of explanation has been understood and elucidated in the experimental non-human primate, such as the macaque (i.e., the rhesus monkey). Nevertheless, in humans, we have not been able yet to determine with certainty precise and specific cortical structural connectivity beyond the stems of the principal fiber pathways (Makris et al., 2023a, 2023b; Rushmore et al., 2020). Thus, the specific origins and terminations of fiber tracts in the human brain are generally undetermined, which provides an incomplete understanding of structural connectivity (e.g., Rushmore et al., 2020). Recent advances in diffusion-based MRI tractography have been hampered by this gap in knowledge of human brain structural connectivity (Makris et al., 2023b; Mesulam et al., 2015; Pandya et al., 2015). Besides that, anatomical understanding, an important factor affecting knowledge of the specific circuitry underlying a cortical area, is hindered by the structural and functional neuroanatomical variability between individuals. Structural and functional variability in humans is a critical factor, especially with personalized medical interventions such as TMS. To minimize these uncertainties, current neuroimaging and neurophysiology technologies can be implemented. More specifically, navigated TMS combined with real-time tractography (Aydogan et al., 2023) helps in detecting the structural cortical and subcortical connectional matrix of the targeted cortical area (Makris et al., 2023a, 2023b). On the other hand, TMS combined with fMRI and high-density electroencephalography (EEG) could be implemented to address the functional signature of the stimulated cortical area (Rosanova et al., 2009b) and its associated functional connectivity. Following this mapping approach, we can optimize the anatomical, connectional, and functional ontologies of a targeted area (Makris et al., 2023a).

### Adjacent circuitries with categorically different behaviors

By extension, we need to characterize the behavioral and clinical ontology of each cortical target in a clinical setting (Makris et al., 2023a). “Silent” regions in the brain, i.e., areas without observable behavioral responses, deserve special consideration, given that they remind us of the existence of a vast cortical territory of uninterpretable brain function. In part, our inability to determine the role of these areas lies in the technical limitations in clinical behavioral assessment. Another reason could be that our tools cannot elicit and/or detect such activities. In the context of a specific pathology, the behavioral clinical ontology of cortical targets in neuromodulation could be addressed based on their neurophysiological signatures (Table 1; Fig. 1) and gathered information from their anatomical and structural connectional architectures (Fig. 2). For instance, in major depressive disorder (MDD), or more specifically in one of its subtypes, the anhedonic one, it may be of great importance to characterize the specificity of the target. Anhedonic MDD is associated with an alteration of the mesocorticolimbic system, which is represented principally by the medial forebrain bundle (MFB) (Yang et al., 2015, 2014). Targeting the MFB at the cortical level requires diffusion MRI-based tractography mapping to identify the different cortical endings of this fiber system. Based on that information, a strategic decision needs to be formulated in planning target engagement. To this end, more advanced brain stimulation techniques need to be implemented, such as real-time tractography and the combined new hdEEG-multi-locus navigated TMS, to ensure specificity of the desired target engagement in terms of a) its accurate cortical location, b) its specific circuitry signature and, c) its specific EEG signature. Fig. 1 illustrates this concept by means of nTMS–EEG (Casarotto et al., 2022; Lioumis and Rosanova, 2022; Rosanova et al., 2009b). As shown in Fig. 2, pre-SMA and SMA may need to be stimulated simultaneously to engage prefrontal, parietal and brainstem structures. Thus, by combining real-time tractography with results from concurrent mTMS and hd-EEG, we could ensure engagement of adjacent cortical areas (pre-SMA and SMA) with categorically different cytoarchitectonic, structural connectivity, and neurophysiological signatures and thus expect to elicit the whole array of beneficial different behavioral and clinical outcomes.

**Figure 2.**
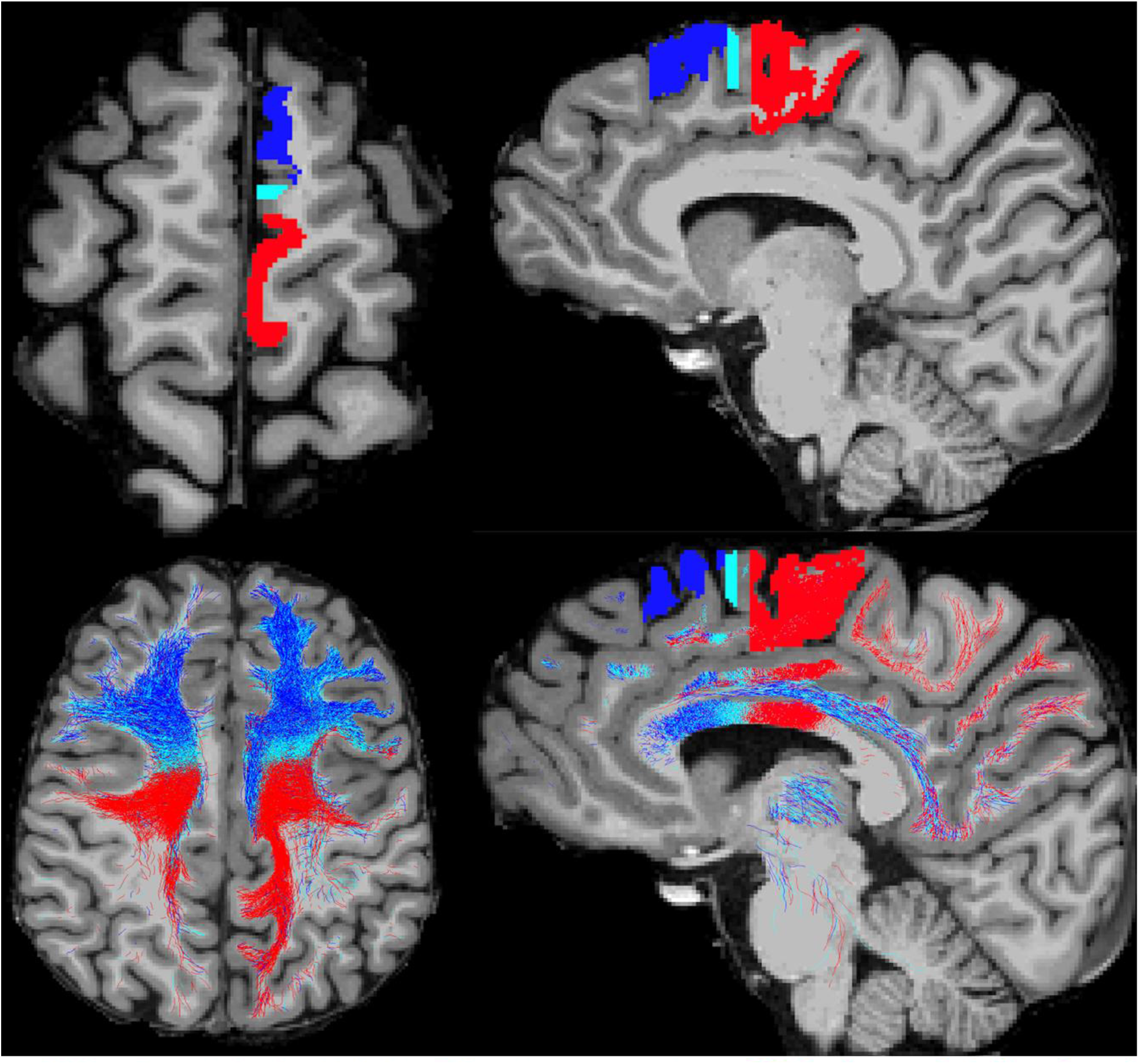
The parallel anterior-posterior gradient in anatomical location and structural circuitry of pre-SMA and supplementary motor area (SMA) in the human brain. The pre-SMA (dark blue), pre-SMA border zone (celestial blue) and SMA (red) are shown in an axial (upper left) and a sagittal (upper right) view. Their structural connectomes as reconstructed by dMRI tractography are shown in an axial (lower left) and a sagittal (lower right) view, following the same color-coding schema, i.e., pre-SMA connectome in dark blue, pre-SMA border zone connectome in celestial blue and SMA connectome in red. The pre-SMA prominent prefrontal circuitry (in blue) located anteriorly, contrasts sharply with the SMA parietal circuitry located posteriorly, whereas the pre-SMA border zone circuitry shows a mixture of both, prefrontal and parietal connections.

## Conceptual and practical consequences

### Integrating structural and neurophysiological signatures in the human cerebral cortex

Technological advancement since the 1950’s allowed us to study the human brain in unprecedented ways. Namely, neuroimaging and neurophysiological techniques enable non-invasive studies of the human brain in vivo. More specifically, anatomical T1-, T2- and diffusion-weighted MRI can provide detailed information about cortical structure and its connectivity. Furthermore, the neurophysiological aspect of the cerebrum can be assessed by fMRI and magnetoencephalography (MEG) in a way that we can generate an understanding of a brain function within space and time resolution parameters at the millimeter and millisecond scales of spatial and temporal resolution, respectively. The integration of structural and functional techniques to novel visualizations of neuronal processing has changed dramatically since the late 1990s, especially with the advent of digital brain-image inflation and other 3D rendering techniques of structural brain data (3D Slicer image computing platform, n.d.; Dale and Sereno, 1993). The latter approaches can be combined with TMS–EEG, which adds further possibilities in studying brain processes related to “causality” (Ilmoniemi et al., 1997; Massimini et al., 2005), a matter of great importance in science, philosophy, and modern neuroscience. Furthermore, this is crucial in elucidating how brain systems break down by disease and how they recover after treatment. Therefore, an emerging concept based on the above consideration is that the elucidation of processing of brain functions, such as language, memory, and affect, can lead to an understanding on how the brain works as an integrated whole and in segregated networks. Ultimately, the anatomical, connectional, functionalistic, and behavioral/clinical ontologies could be integrated in the setting of a “how the brain works” hub.

### Pre-treatment targeting preparation for specific therapeutic interventions

Guiding multi-locus TMS (Nieminen et al., 2022) based on anatomical MRI, real-time dMRI tractography, and real-time TMS–EEG mapping (Fig. 3), as demonstrated manually with compatible nTMS in the present study in one healthy subject, enables us to precisely identify a target cortical area, its underlying circuitry and its neurophysiological signature, semi-automatically or fully automatically. In practical terms, these neuroimaging and neurophysiological procedures need to be applied prior to TMS intervention to ensure reliable target engagement and the optimal TMS intensity dosage in the individual subject. Although in principle obvious, this concept has not been clinically practiced to date. Generally, what is used in current standard-of-care TMS therapeutics is a universal targeting procedure based on standard *x*,*y*,*z* MNI or Tailarach coordinates (for all subjects), which do not necessarily represent accurately the individual subject’s cortical anatomy. However, the TMS stimulation procedure outside the motor cortex (e.g., dorsolateral prefrontal cortex) entails specific coil orientation with respect to the morphological correlates of the primary motor cortex (i.e., central sulcus).

**Figure 3.**
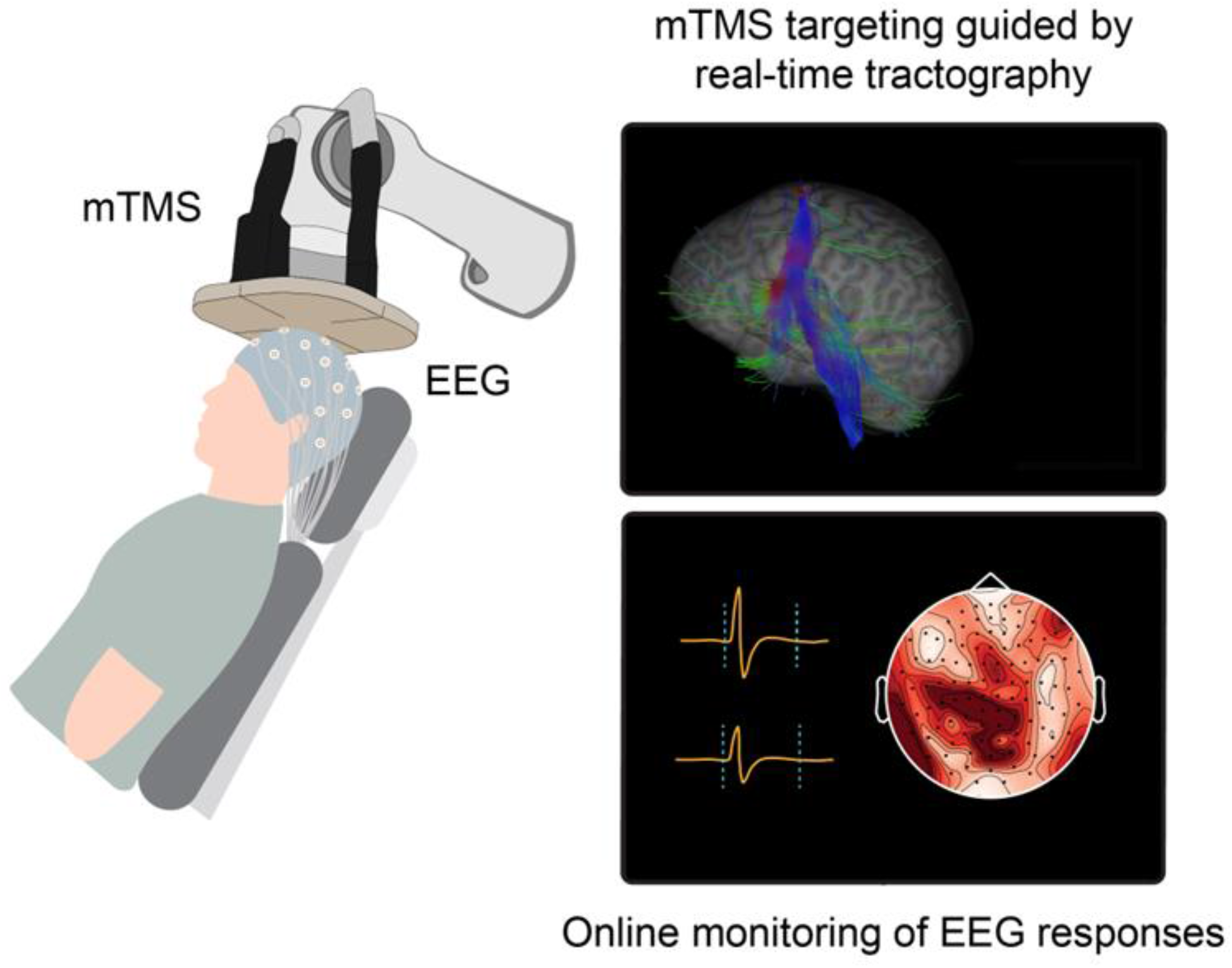
Real-time tractography- and on-line EEG-guided multi-locus TMS.

Consequently, target identification, engaged circuitry and neurophysiological outcome of the target area can be anatomically inaccurate. Therefore, to engage accurately the desired cortical targets, the above-mentioned neuroimaging procedures need to be performed prior to the therapeutic intervention. Thus, processing of multimodal neuroimaging data could enable the precise anatomical determination of the cortical target and ensure more specific and reliable treatment planning. Conceptually, structural and electrophysiological specificity of cortical target and circuitry and their reliable engagement is currently feasible following the methodology discussed herein.

